# Nucleoside antibiochemotherapy repressed the growth, chemoresistance, survival, and metastatic potentials of castration-resistant prostate cancer cells

**DOI:** 10.1101/2021.08.22.457225

**Authors:** Saheed Oluwasina Oseni, Genesis Acosta Laguer, Faika Ambrin, Magdalah Philemy, Javoncia Betty, James Kumi-Diaka

**Author notes:** **CORRESPONDING AUTHORS:** Saheed Oluwasina Oseni, *DVM, MS, Ph.D.*, James Kumi-Diaka, *DVM, MS, Ph.D, DSc.*, Florida Atlantic University, 3200 College Avenue, Davie, FL 33314.

## Abstract

There is currently no definitive cure for metastatic castration-resistant prostate cancer (mCRPC), therefore justifying the incessant need for more investigative studies to either repurpose old drugs or identify novel and effective therapeutics. In this study, we investigated the possible anticancer effects of two nucleoside antibiotics: puromycin and blasticidin. We hypothesized that the two antibiotics alone or combined with other drugs will inhibit prostate cancer (PCa) cell proliferation and metastasis and induce cell death via apoptosis. mCRPC cell lines (PC3 and DU145) with different p53-gene statuses were cultured and seeded in 96 well-plates, and thereafter treated with varying concentrations of puromycin and blasticidin (1 ng/mL - 100 μg/mL) for 24 - 48 hours. Resazurin reduction and/or MTT assays were done to evaluate the treatment-induced effects on mCRPC cell viability and proliferation. The colony-forming assay measured the cell survival rate following treatment nucleoside antibiotics while scratch migration assay and dual-fluorescent microscopy assessed the effects on metastatic potential and cell death, respectively. The two antibiotics were combined with either paclitaxel, docetaxel, or cabazitaxel to check for synergism. Our results indicate that both antibiotics exhibit dose- and time-dependent anticancer effects on growth, survival, and metastasis of mCRPCs. PC3 cells were significantly more susceptible to both antibiotics compared to DU145 cells. Both cell lines were more susceptible to puromycin compared to blasticidin. Synergism was observed when each antibiotic compound was combined with any of the three taxanes. In conclusion, we have demonstrated that both puromycin and blasticidin could be explored for the treatment of mCRPC.

**GRAPHICAL ABSTRACT:** 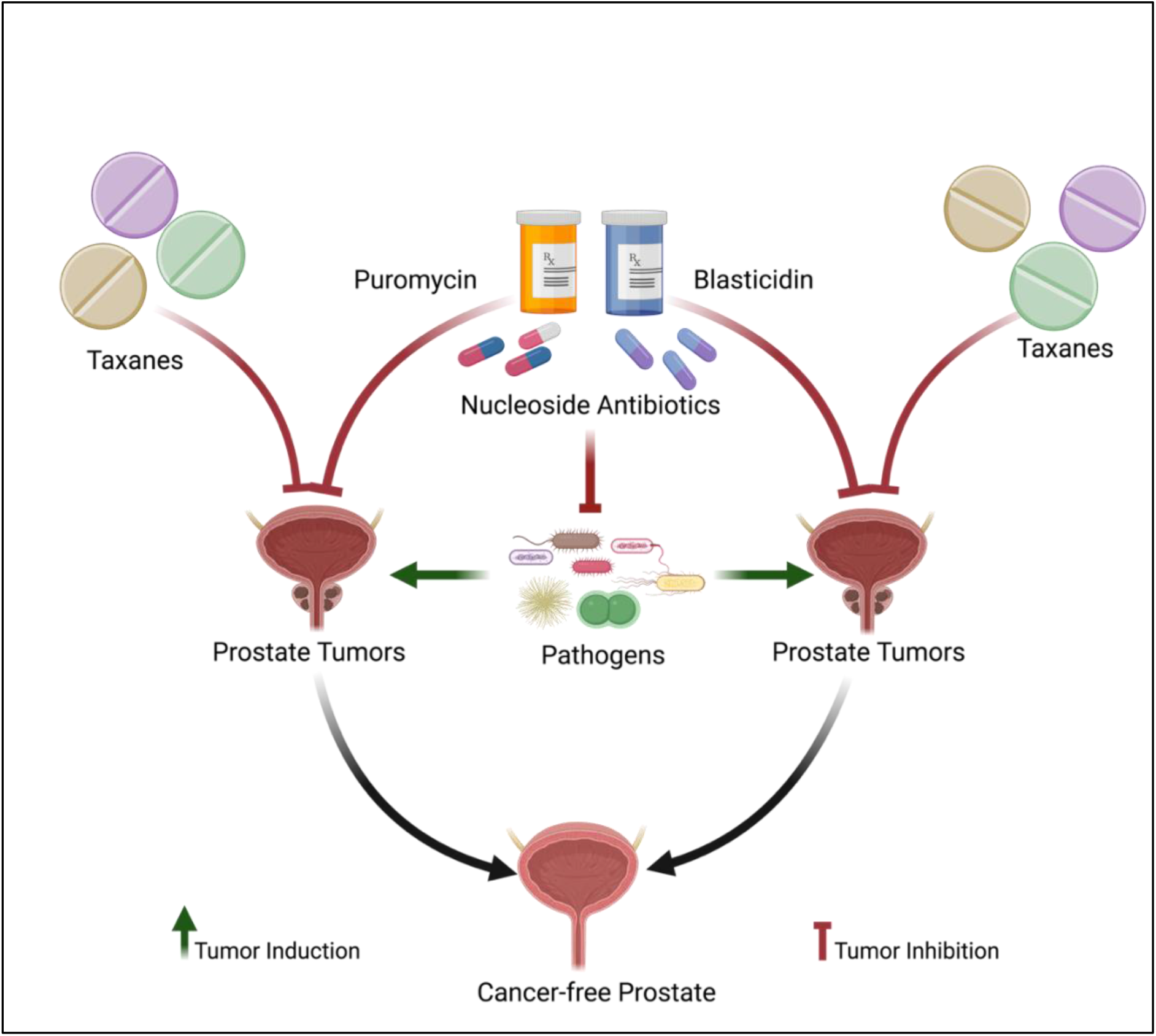

## INTRODUCTION

Prostate cancer (PCa) currently affects more than 2 million men in the United States. Estimates have it that for the year 2021, about 248,530 new cases and about 34,130 deaths due to PCa are expected [**1**]. Approximately one in forty-one men in the United States are expected to die from PCa [**2**]. PCa can develop as adenocarcinomas, neuroendocrine tumors, small cell carcinomas, or transitional cell carcinomas, however, the most common form is adenocarcinoma, which is an abnormal growth of the secretory cells of the prostate gland [**3, 4**]. The severity of PCa can vary, most prostate adenocarcinomas are localized with gradual growth and recognized to be less aggressive forms of PCa [**5**]. In more severe cases, prostate tumors can grow rapidly and metastasize to different areas of the body [**6**]. The less aggressive forms of PCa have a 90% five-year survival rate and can be treated by radiation therapy, surgery, and androgen deprivation therapy (ADT); the latter focuses on reducing the availability of androgens to PCa cells. Whereas, as soon as PCa becomes castration-resistant and metastatic, the survival rate drops to around 30% [**7**]. The aggressive forms of PCa can be treated with chemotherapy which reduces cancerous cells but also diminishes the quality of health [**8**]. The advanced or aggressive stage of the disease remains incurable, hence justifying the demand for more effective therapeutic strategies, including the repurposing of older drugs to effectively treat metastatic castration-resistant PCa (mCRPC).

Though most antibiotics are administered for antimicrobial purposes, a few others have been shown to also have cytotoxic effects on mammalian cells [**9, 10**]. For instance, anthracycline antibiotics such as daunorubicin, doxorubicin, epirubicin, idarubicin, and valrubicin, as well as non-anthracycline antibiotics such as bleomycin, dactinomycin, mitomycin-C, and mitoxantrone exhibit antitumor activities [**11, 12**]. Puromycin and blasticidin are examples of nucleoside antibiotics with verified cell killing activities on both prokaryotic and eukaryotic cells [**13, 14**].

Puromycin (3′-[α-Amino-*p*-methoxyhydrocinnamamido]-3′-deoxy-N, N-dimethyladenosine dihydrochloride) is an amino-nucleoside antibiotic, isolated from *Streptomyces alboniger* bacteria [**13**]. Puromycin is known to kill prokaryotic and eukaryotic cells by causing premature chain termination during protein translation [**15**]. Some studies have shown that puromycin can induce cell death in leukemia and ER-negative breast cancer [**15**]. The most established mechanism of action of puromycin involves the 3’ terminal end of aminoacyl-tRNA in which the antibiotic molecule binds or inserts itself into a growing polypeptide chain resulting in premature termination [**16**]. This termination consequently inhibits protein synthesis. Thus leading to rapid cell deaths at effective (microgram) antibiotic concentrations [**15**].

Blasticidin-S (3-amino-5-[carbamimidoyl(methyl)amino]pentanoyl]amino]-6-(4-amino-2-oxopyrimidin-1-yl)-3,6-dihydro-2*H*-pyran-2-carboxylic acid) is a peptidyl-nucleoside, initially isolated from *Streptomyces griseochromogenes* in 1958 [**13**]. However, other Streptomyces species such as *Streptomyces setoni, Streptomyces griseoflavus, Streptomyces albus subsp. pathocidicus*, and Streptoverticillium sp. JCM 4673.4, has been shown to also produce blasticidin. Mechanistically, blasticidin acts on both eukaryotic and prokaryotic cells through the inhibition of the termination step of protein synthesis which reduces peptide bond formation [**17**]. Unlike some broad-spectrum antibiotics such as chloramphenicol and linezolid that bind to the A site of the large subunit of ribosomes, blasticidin binds to the P site of the ribosomal unit where the peptidyl-transferase is being inhibited [**18**]. It has been shown that blasticidin competes with the A-site, hence binding the same way as puromycin [**19**].

However, there is the possibility that mammalian cells that have a resistance to blasticidin may also develop some resistance to puromycin, thus suggesting that the therapeutic potential of blasticidin and puromycin could be inhibited by blocking the influx and binding of these drugs to the A-site [**20**]. Furthermore, the use of blasticidin as a selective agent during genetic engineering was justified with the discovery that mammalian cells with no resistant genes are antibiosensitive or die when exposed to blasticidin concentrations of 1 µg/mL - 10 µg/mL while bacteria cells were antibiosensitive or die at a dose of 25 µg/mL - 100 µg/mL at approximately 3 - 7 days post-exposure; toxicity (LD_50_) in mice was at 2.8 mg/kg [**21**]. Blasticidin has also been suggested in a few studies to have antifungal, antitumor, antiviral, antibacterial, anti-inflammatory properties [**22**].

It is worth noting that not many studies are available in the literature with regards to the anticancer potential of nucleoside antibiotics, especially at low doses [**23**]. We, therefore, hypothesized that both puromycin and blasticidin antibiotics alone or in combination with other anticancer drugs will inhibit PCa cell proliferation, metastasis, and induce cell death via apoptosis, in a dose- and time-dependent manner. To confirm our hypothesis, we investigated the anticancer effects of puromycin and blasticidin antibiochemotherapy on mCRPCs to assess if these antibiotics can inhibit PCa growth and metastatic potential (cell migration).

## MATERIALS AND METHODS

### Cell Culture and Growth Conditions

In this study, two castration-resistant PCa (CRPCs) cell lines were used. DU145 (ATCC HTB-81) and PC3 (ATCC CRL-1435) PCa cell lines were purchased from American Type Culture Collection (ATCC; Manassas, VA). The PC3 cell line is an androgen-insensitive PCa that was derived from the bone metastasis of grade IV prostatic adenocarcinoma of a Caucasian patient whereas DU145 is an androgen-insensitive PCa, isolated from the brain metastasis of grade II Caucasian PCa patient. PC3 cells are adherent mCRPCs with a doubling time of 24 hours while DU145 cells are also adherent mCRPCs with a doubling time of 34 hours. PC3 cells are known to have a higher metastatic potential than DU145 cells. Both cell lines were maintained in tissue culture flasks containing RPMI1640 complete growth media supplemented with 10% fetal bovine serum (Biological Industries) in an incubator at 37°C and 5% CO_2_; until they became ∼80% confluent in about 2 - 3 days. We avoided supplementing the complete media with penicillin-streptomycin antibiotics (1%) to prevent any interference with our antibiochemotherapy experiments.

### Drug Preparation and Reconstitution

Puromycin (10 mg/mL) and blasticidin (10 mg/mL) were purchased from Fisher Scientific (Waltham, MA; **Figure 1**). Stock solutions (100 µg/mL) were prepared and further reconstituted to working solutions at graded concentrations ranging from 1 ng/mL – 100 µg/mL. The remaining stock solutions were stored at -20°C until needed. Paclitaxel (#A4393), docetaxel (#A4394), and cabazitaxel (#B2157) were purchased from APExBio (Houston, TX) at a stock concentration of 10 mM in 1 mL DMSO. The stock solution of each compound was reconstituted into working concentrations: paclitaxel (0, 0.01, 0.1, 1.0, 10 μM), docetaxel (0, 0.01, 0.1, 1.0, 10 μM), and cabazitaxel (0, 0.01, 0.1, 1.0, 10 μM) in anticipation for combination treatment experiments.

**Figure 1:**
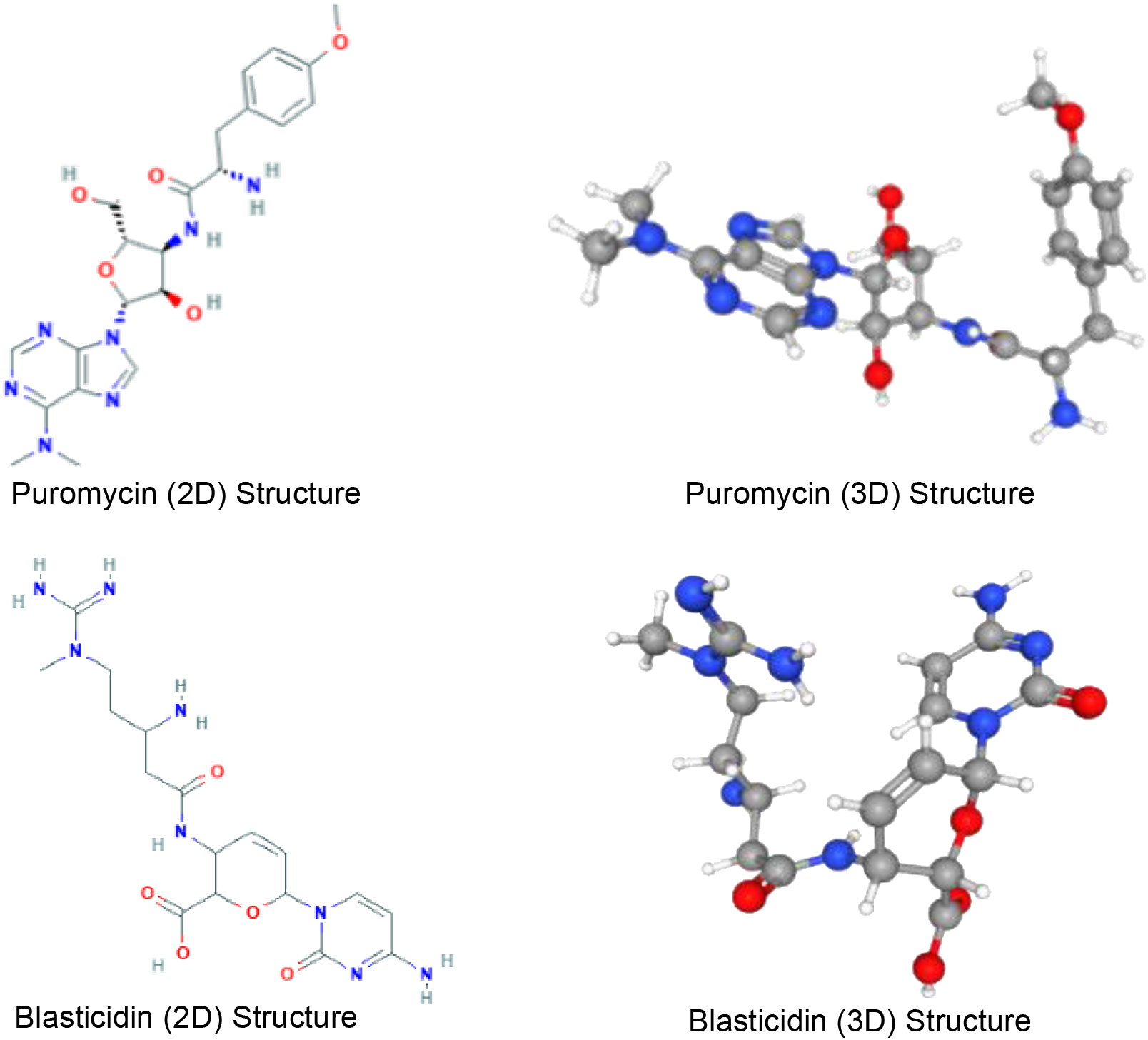
The 2D and 3D chemical structures of puromycin and blasticidin antibiotics. Adapted from https://pubchem.ncbi.nlm.nih.gov/

### Cell Plating and Dosing with Antibiotics

Using the cultured DU145 and PC3 cell lines, viable PCa cells were counted per ml using Trypan blue exclusion assay. Using a trypan blue solution, a dilution factor of 2 (i.e., 10 µL of the cell suspension to 20 µL of trypan blue and placing the mixture onto a hemocytometer) was made. Then, the cells were observed by mounting the hemocytometer on the microscope stage and viewing the cells through the lens of the microscope. The cells were counted using a cell counter. To calculate the total amount of PCa cells per mL, the average viable cells counted had to be multiplied by 10,000 and a dilution factor of 2. 10^4^ cells were seeded into each well of the 96-well plates and incubated for 24 hours for the attachment to occur. Thereafter, the cells were exposed to 100 µL of graded pre-determined (serial) concentrations of antibiotics (1 ng/mL - 100 µg/mL) and incubated for 24 - 48 hours at 37°C and 5% CO_2_. For the combination treatment experiment, 10^4^ PC3 cells were seeded into each well of the 96-well plates and incubated for 24 hours before treatment. Thereafter, the cells were pre-exposed to IC_50_ of either puromycin or blasticidin antibiotics and thereafter treated with predetermined IC_50_ of paclitaxel, docetaxel, and cabazitaxel, followed by incubation for 24 - 48 hours at 37°C and 5% CO_2_ before assaying.

### Cell Viability and Proliferation Rate Assessment

The DU-145 and PC3 cells were cultured until they reached 90% confluency and 10^4^ of cells were seeded into 96-well plates. Then, they were exposed to (1 ng/mL - 100 µg/mL) blasticidin and puromycin in which one row of each plate was left unexposed to antibiotics and/or treated with 1% DMSO (vehicle) to serve as a control. The plates were incubated at 37°C and 5% CO_2_ for 48 hours. From there, the MTT and Resazurin reduction assays were used to observe the cell viability and proliferative rate. This was done by adding 10 µL of MTT solution to the appropriate wells or 20 µL of Resazurin solution to the proper wells and incubated for 4 hours at 37°C and 5% CO_2_. Once the MTT plates were done incubating, 100 µL of DMSO was added to further break down any formazan crystallizations. Finally, the spectrophotometer (BioTek) was used to quantitatively analyze and obtain the absorbance values for each well. For the MTT plates; the spectrophotometer had to be calibrated to 570 nm, and for the Resazurin plates, it needed to be 560 nm to record the proper absorbance. The MTT and Resazurin reduction assays use color change to indicate the efficiency of the antibiotics by indicating any reduction of cell proliferation; the richness of the color has a direct relationship with the number of viable cells. When using the MTT assay, cells that are recognized as viable and having an active metabolism can convert MTT (yellow) to formazan resulting in purple color. When using the resazurin assay, viable cells with an active metabolism are indicated by a change to pink color in which Resazurin is converted to Resorufin.

### Cell Death Assay and Dual Fluorescent Microscopy

The Acridine orange (AO) and propidium iodide (PI) dual-staining assay was used to determine the mode of treatment-induced cell death for both PC3 and DU145 PCa cells. This assay was used to differentiate between live, early apoptotic, late apoptotic, and necrotic PCa cells. For this experiment, 8-well chamber culture slides were purchased (SPL Life Sciences, Korea) and used to culture 10^4^ cells per well. After attachment, cells were exposed to graded concentrations of puromycin (1 ng/mL - 10 µg/mL) and blasticidin (1 ng/mL - 10 µg/mL) and then incubated at 37°C and 5% CO_2_ for 48 hours. A cocktail (1:1) of 1 µL of 0.5 mg/mL acridine orange and propidium iodide (AO/PI) dyes in PBS were created and added to each well of the culture slides for 10 minutes at room temperature and thereafter rinsed in water to remove excess dyes. AO dye is known to enter live cells as a green color fluorochrome, whereas PI can only enter if the cell wall is damaged. Cells that emit a yellow/ brown/red color fluorescence indicate early and late apoptosis, respectively. Dual-fluorescence was measured using a fluorescent microscope (Nikon Eclipse E600) with an excitation wavelength of 460 nm (green/FITC channel) and an emission wavelength of 650 nm for AO and an excitation wavelength of 525 nm (red channel) and an emission wavelength of 595 nm for PI.

### Colony Formation Assay

mCRPCs are known to exhibit aggressive and anchorage-independent cell growth and high clonogenic potential. In this study, we used the colony formation assay to determine the treatment-induced effect of both antibiotics on the colony-forming ability of PC3 and DU145 PCa cell lines. 10^3^ cells (DU145 and PC3) were separately seeded into each well of 12-well plates and treated immediately with a graded concentration of puromycin (1 ng/mL - 10 µg/mL) and blasticidin (1 ng/mL - 10 µg/mL). Antibiotics were removed after 24 hours. We then plate the PCa cells and incubate them at 37°C and 5% CO_2_ in a humidified incubator for 7-days. Thereafter, the PCa cells were stained using crystal violet dye. This histological stain helps to count and identify all the colonies that were formed under the microscope. This technique of colony formation is an *in vitro* cell survival assay that helps us to identify single tumorigenic cells that are resistant to antibiotics with the ability to create colonies. Also, this bioassay helps to determine the dose-dependent efficacy of the nucleoside antibiotics in inhibiting the growth and survival of PCa. Colonies were visualized and counted with the aid of a Stereomicroscope (Zeiss AX100). Image analysis and visualization were performed using Image J software to derive colony confluency indexes (%) for each treatment group.

### Scratch Migration Assay

Scratch migration assay can be used to determine the metastatic potential of PCa cells from one part of the body to another, mimicking cell migration. About 10^4^ of DU145 and PC3 cells were plated in a 12-well plate and allowed to become confluent and attach. Following a 100% confluency of cells, a straight-line scratch was made at the center of each well of each plate containing both PC3 and DU145 cells. Thereafter, cells were then treated immediately with a graded concentration of puromycin (1 ng/mL - 10 µg/mL) and blasticidin (1 ng/mL - 10 µg/mL). Images of the scratches were analyzed using ImageJ software. The scratch diameters were compared between control, pre-treatment, and post-treatment (48 hours) wells to determine the antimetastatic activity of puromycin and blasticidin. Image analysis and visualization were performed using Image J software to derive migration indexes (%) for each treatment group.

### Statistical Analysis

Data representation is expressed by means ± standard deviation (SD) in quadruplicates. Each experiment was repeated independently three times to confirm and validate the results. Student’s t-test as well as one-way and two-way ANOVA were used to test our hypothesis and the significance of differences between mean results and the control. The p-value of < 0.05 was accepted as significant. The concentrations of either the antibiotic or taxane alone or in combination that inhibits 50% of cell proliferation (IC_50_) were determined by fitting the dose-response curves utilizing the non-linear regression model in GraphPad Prism software (version 9.0). Both the dose-reduction index (DRI) and combination index (CI) were calculated as described by Chou and Talalay [**24**].

## RESULTS

### Puromycin and blasticidin inhibit the proliferation and cell viability of mCRPC cells

To determine the inhibitory effects of puromycin and blasticidin on mCRPC cells, both MTT and resazurin assays were performed which allowed us to assess the ability of both antibiotics to reduce viability and proliferation of mCRPCs compared to control or untreated cells (**Figure 2**). After 48 hours of treatment, the dose-dependent effect of each antibiotic was evaluated using cell viability assays. **Figure 2** plot describes the dose-response effects in which a negative correlation between the concentration (dose) of antibiotics and cell viability in both cell lines was observed. At the antibiotic concentrations ranging from 1 ng/mL to 100 µg/mL, the cell viability was found to be the highest for 1 ng/mL and lowest at 100 µg/mL. In PC3 cells treated with puromycin and blasticidin, we observed approximately 0% cell viability from around 25 µg/mL upward, however, for DU145, 0% cell viability was observed when treated with 100 µg/mL after 48 hours. After MTT assay and statistical analysis, puromycin and blasticidin were found to have lower IC_50_ in PC3 cells compared to DU145 cells (**Figure 3**). This means that puromycin and blasticidin inhibit 50% of PC3 cells at a lower concentration compared to DU145 cells. The concentration of antibiotics required to inhibit 50% of mCRPCs (IC_50_) are: Puro-PC3 (0.48 µg/mL; 95% Confidence Interval: 0.34 – 0.66) < Blast-PC3 (1.17 µg/mL; 95% Confidence Interval: 0.99 – 1.37) < Puro-DU145 (3.25 µg/mL; 95% Confidence Interval: 2.40 – 4.39) < Blast-DU145 (7.65 µg/mL; 95% Confidence Interval: 5.91 – 9.89). Morphologically, PC3 and DU145 cells were observed to show a dose-dependent decrease in their population with most cells exhibiting signs/hallmarks of apoptosis at 48 hours after treatment with puromycin and blasticidin (**Figure 4**).

**Figure 2:**
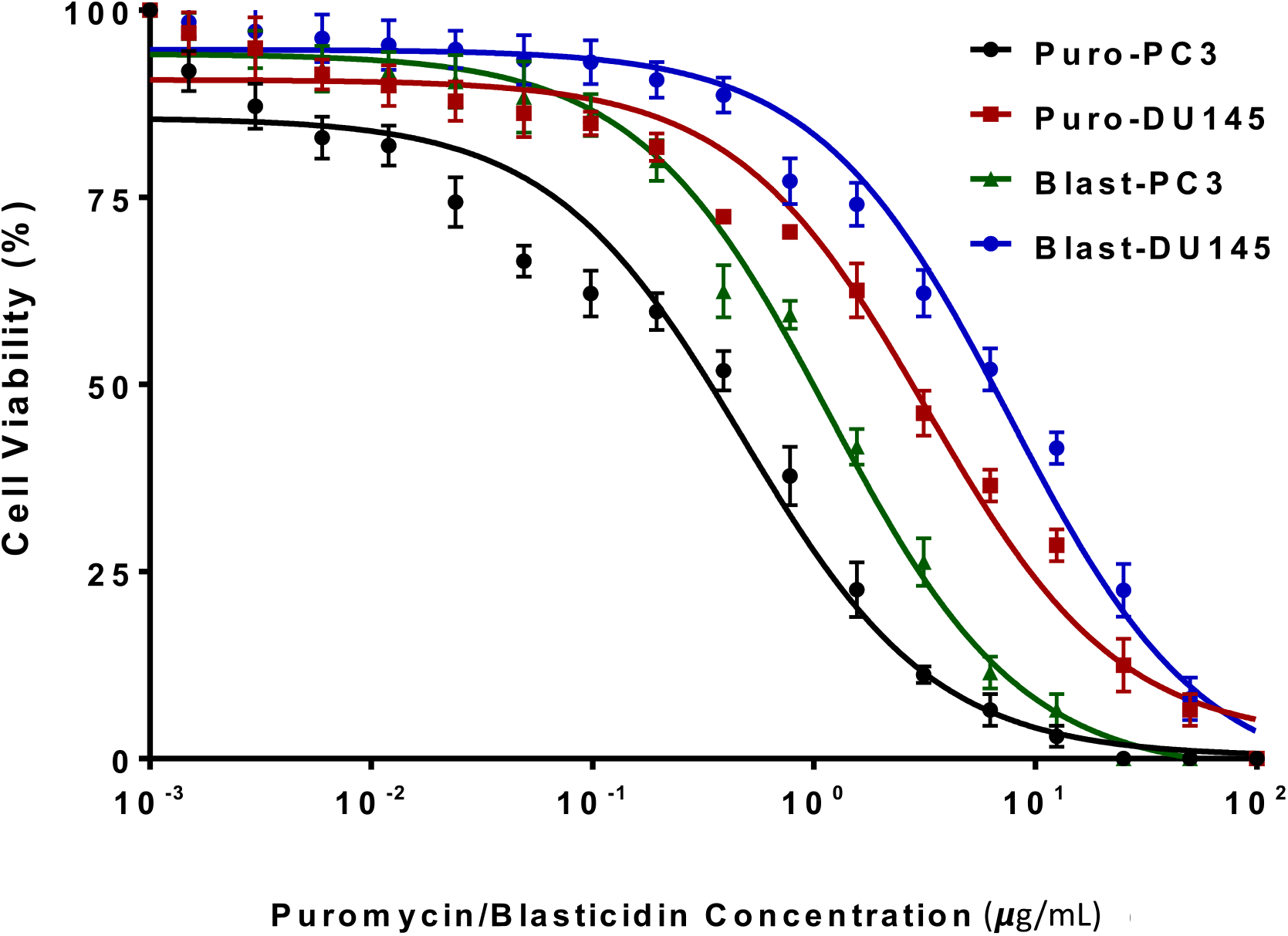
Dose-response plot (non-linear regression fit) of the cell viability of DU145 and PC3 cells after treatment with a graded concentration of puromycin and blasticidin antibiotics for 48 hours. Data are represented as mean ± SD in quadruplicates, in which we repeated the experiment independent 3 times at low concentrations (1 ng/mL – 100 µg/mL). P < 0.05 is considered as significant. We observed a dose-dependent, antibiotics/drug-dependent, and cell phenotype-dependent response with regards to cell viability.

**Figure 3:**
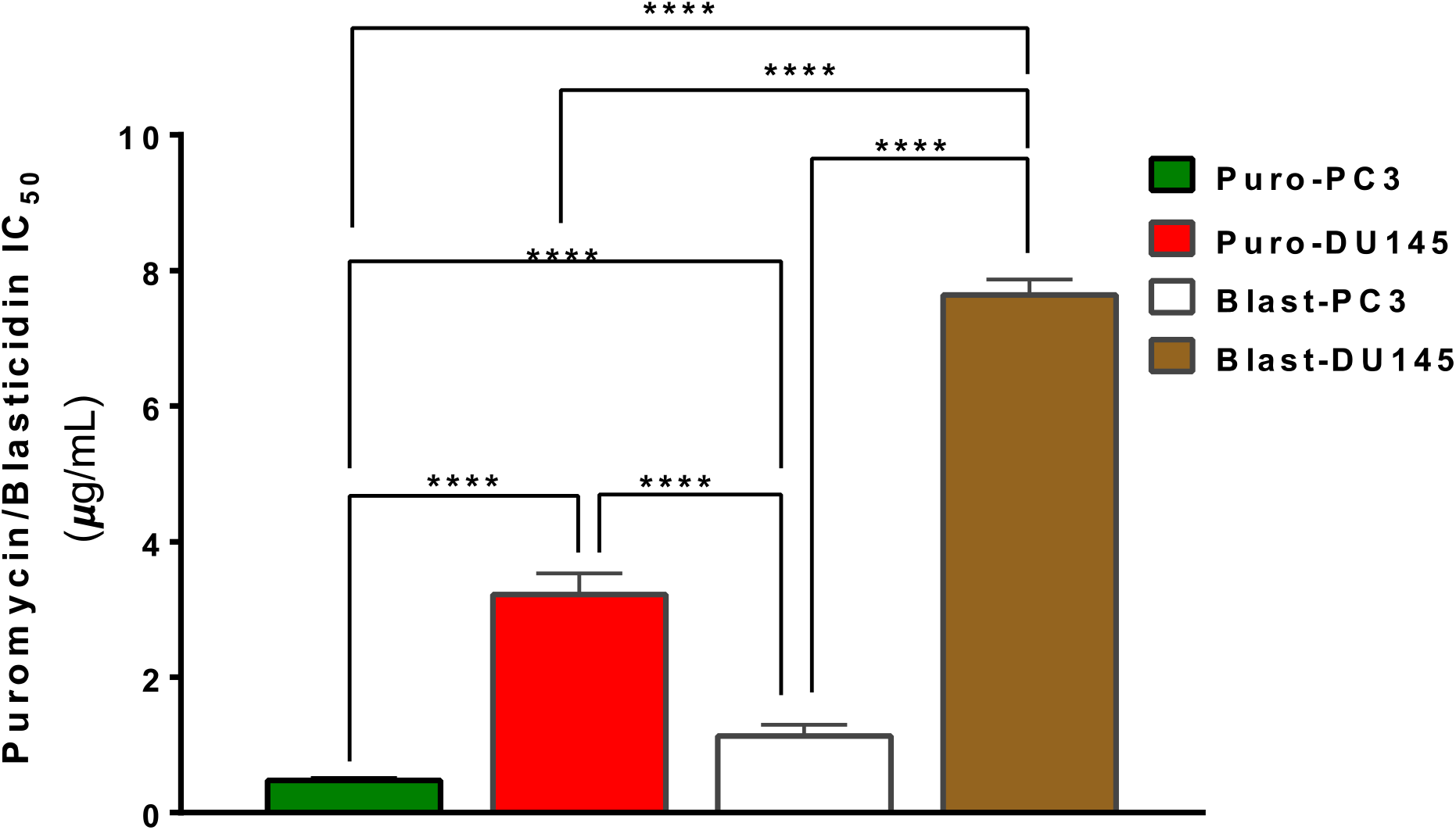
Bar chart plot of dose versus IC_50_ for puromycin and blasticidin in both DU145 and PC3 PCa cell lines. In this figure, we measure the effective concentration that kills 50% of cells. As seen, the Blast-PC3 (Blasticidin-treated PC3) cell line requires a higher concentration to inhibit the cells, whereas the Puro-PC3 (Puromycin-treated PC3) cell line requires the least concentration for the same effect. Blast means blasticidin and Puro means puromycin. The differential effects on cell viability for each drug (IC_50_) per cell line observed are as follows: Puro-PC3 (0.47 µg/mL) > Blast-PC3 (1.17 µg/mL) > Puro-DU145 (3.25 µg/mL) > Blast-DU145 (7.65 µg/mL). *p < 0.05, **p < 0.01, ***p < 0.001, ****p < 0.0001 indicate significant p-values while ns indicates non-significant p-values. Treatments were done in triplicates and repeated at least 3 times independently.

**Figure 4:**
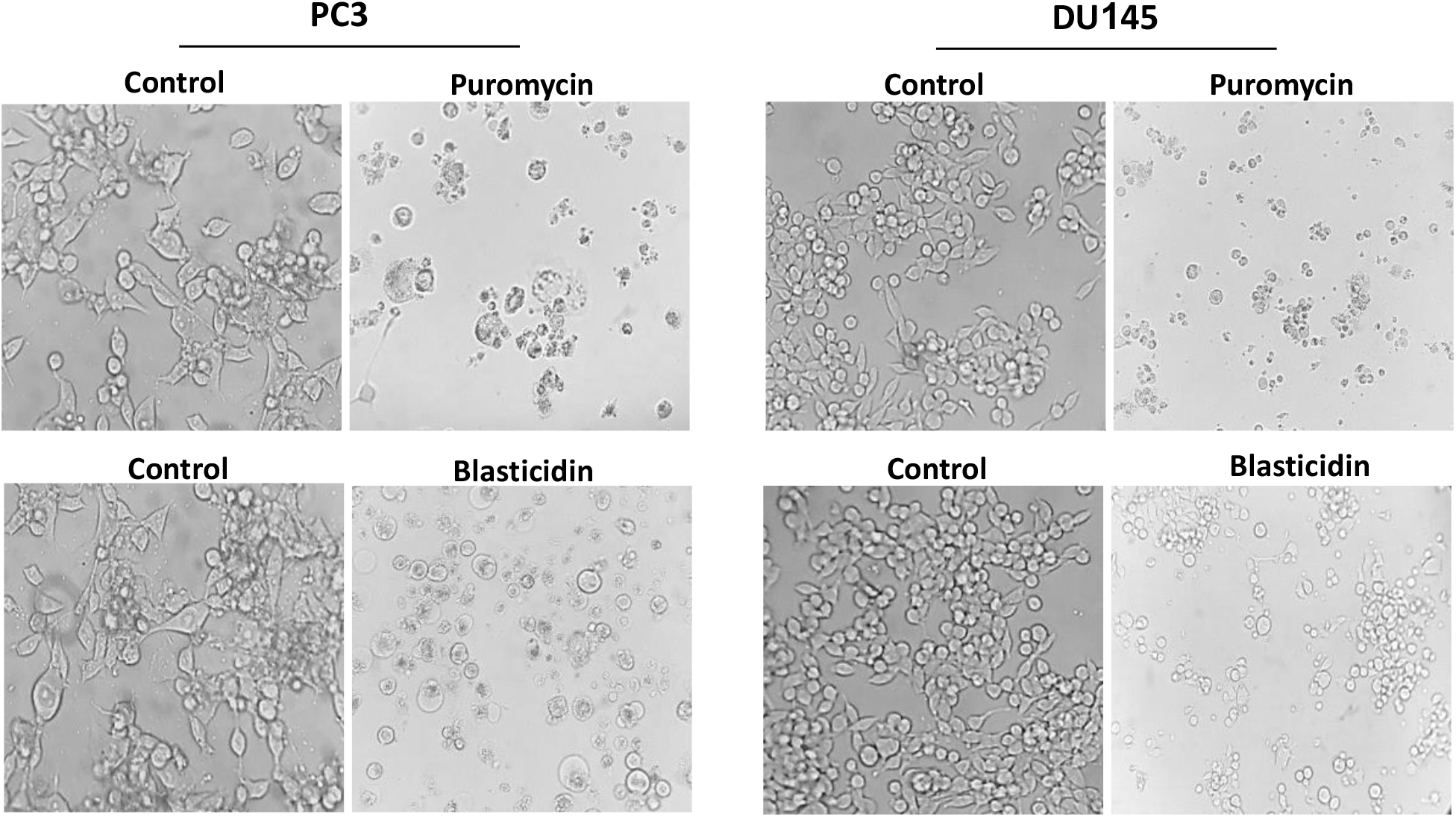
Phase-contrast photomicrographs of morphological and population changes in PC3 and DU145 cells following treatment with puromycin and blasticidin antibiotics relative to control. Microscopic images showing morphological changes in PC3 and DU145 cells in response to antibiotics treatment (IC_50_) compared to untreated control. We observed treated cells showing signs of cell cycle arrest and apoptosis.

### Puromycin and blasticidin antibiotics decreased the cell survival and tumorigenic potential of mCRPC cells

The colony formation assay was used to assess the ability of the antibiotics to inhibit mCRPC colonies, chemoresistance, and the extent to which they did. Since chemoresistance is a method of survival for cancer cells, and the number of colonies depicts the degree of resistance to drugs; more colonies show higher resistance and vice versa. As observed in **Figures 5A & 5B**, at 7 days post-treatment, PC3 cells have little to no colonies left, which indicates that they are more susceptible to both antibiotics compared to the control as well as similarly-treated DU145 cells. In other words, this means that PC3 cells were less resistant to puromycin compared to DU145 cells, even though both still experienced marked cell death. On the other hand, DU145 cells were relatively chemoresistant to blasticidin compared to PC3 cells, as there were more colonies present post-treatment (**Figure 5A**). In our study, the results of the colony formation assay revealed the ability of puromycin, and to a lesser extent blasticidin, to effectively inhibit CRPC survival and chemoresistance (**Figure 5B**).

**Figure 5A:**
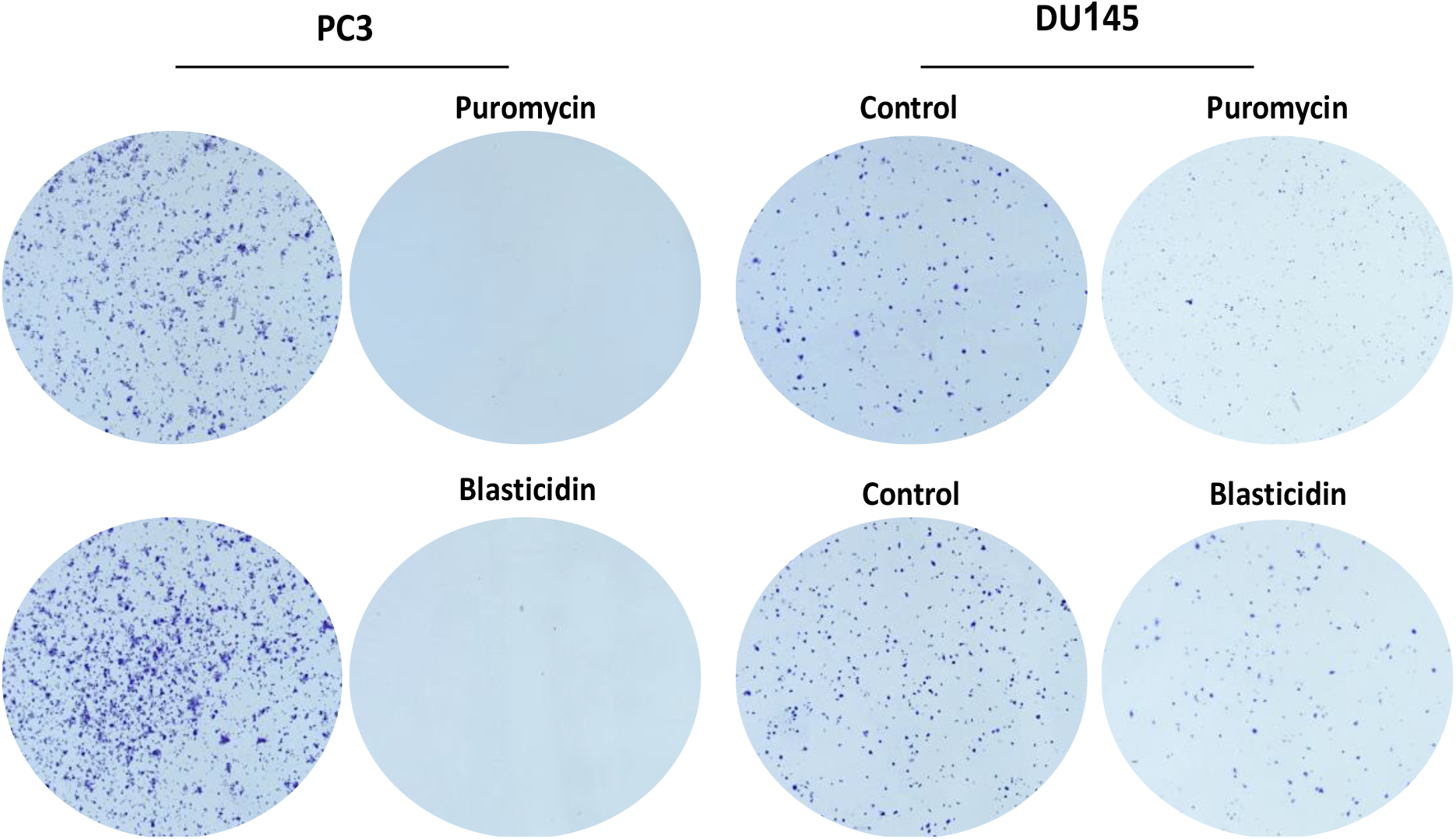
Photomicrograph of PC3 and DU145 cell colonies following treatment with puromycin and blasticidin. Microscopic images comparing the amount of PC3 and DU145 colonies formed in response to treatment with IC_50_ of each antibiotic after 7 days incubation relative to the untreated/control groups. We observed treated cells showing signs of reduced colony formation, indicating an inability to form microtumors.

**Figure 5B:**
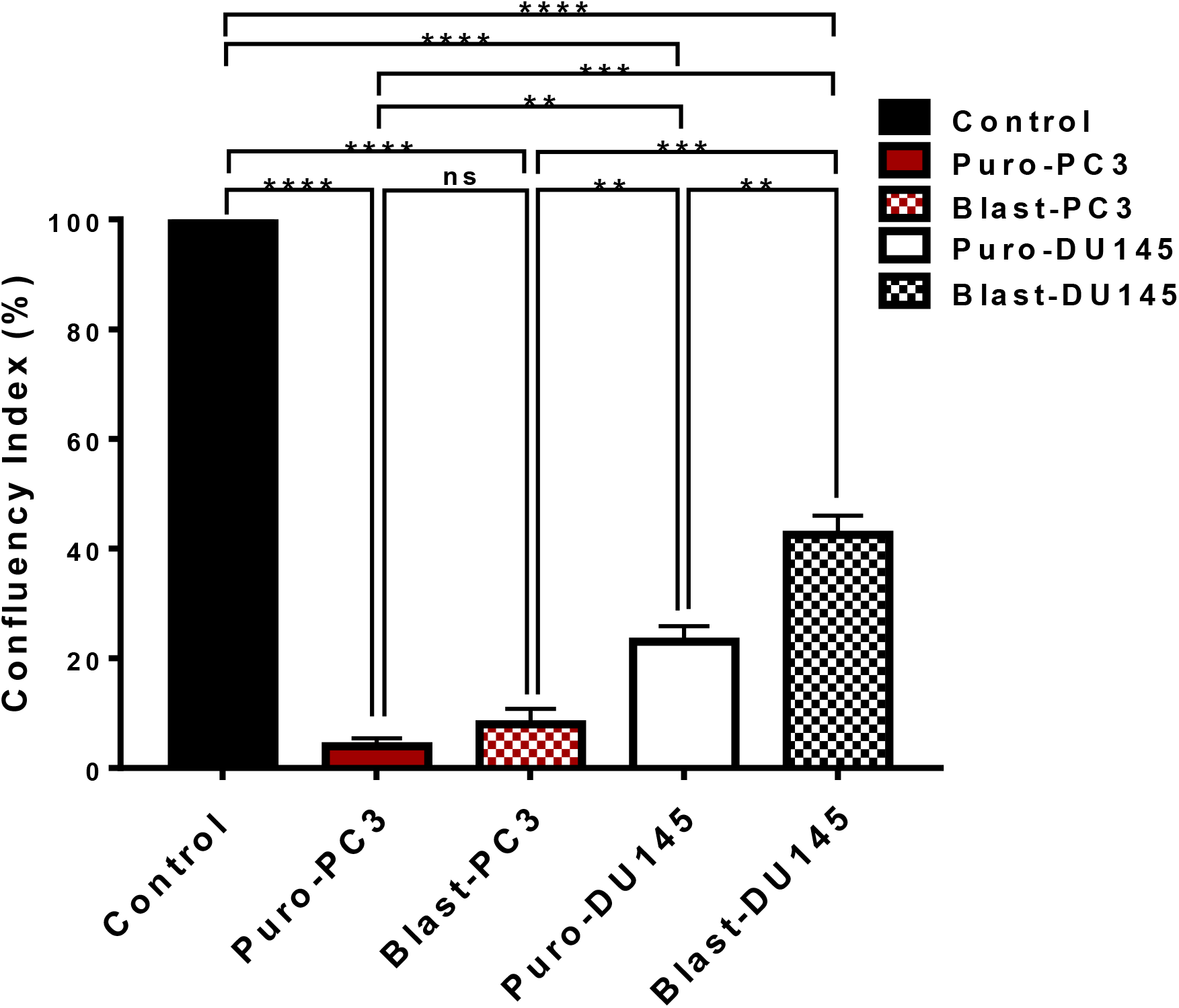
Bar chart comparing colony confluency indexes between puromycin-treated (IC_50_) and blasticidin-treated (IC_50_) PC3 and DU145 cells relative to the control. The wells with colonies were analyzed using Image J to generate normalized numeric values corresponding to the number of individual colonies formed (confluency) after treatment relative to control. The colony confluency index (%) was plotted and compared. *p < 0.05, **p < 0.01, ***p < 0.001, ****p < 0.0001 indicate significant p-values while ns indicates non-significant p-values. Treatments were done in triplicates and repeated at least 3 times independently.

### Puromycin and blasticidin decreased the metastatic potential of mCRPC cells

The ability of puromycin and blasticidin S antibiotics to reduce the metastatic potential of PC3 and DU145 was tested by performing the scratch migration assay (**Figures 6A & 6B**). The untreated PC3 and DU145 cells served as the control, which allowed us to measure the metastatic growth potential of the PC3 and DU145 cell lines in the absence of the antibiotics. For the untreated control cells, we observed a complete closure of the scratches created at about 48 hours post-treatment in both cell lines. However, in the PC3 cell line at 48 hours post-treatment with puromycin or blasticidin, we observed that both inhibit cell migration (metastatic potential) since the scratches created were still visible. The dots at the center of the scratches (**Figure 6A & 6B**) in the treated group represent migrating cells, meaning that migration did occur but at a lower rate when compared to the control. Looking further, the puromycin was more effective than blasticidin in reducing the metastatic potential for PC3 cells because one can observe that there was a higher amount of cell migration at 48 hours post-treatment blasticidin than that of puromycin (**Figure 6C**). The DU145 cell line followed the same trend in which both antibiotics displayed the ability to reduce metastatic potential but puromycin proved to be a more effective antibiotic (**Figure 6C**).

**Figure 6A:**
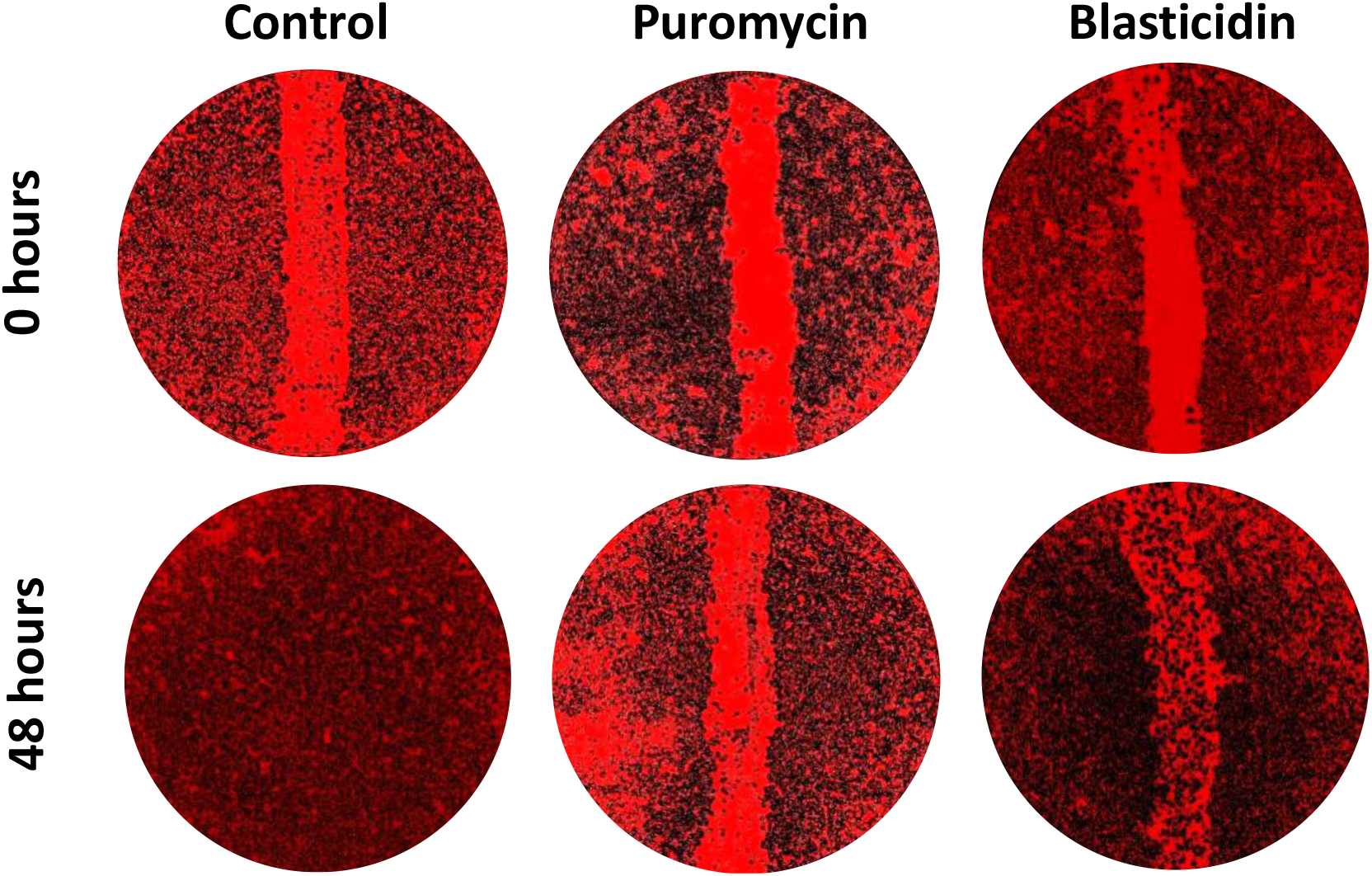
Photomicrograph of scratch cell-migration assay of PC3 colonies following treatment with puromycin and blasticidin antibiotics. Microscopic images (scratch cell-migration assay) showing colonies of PC3 before (0 hours) and after treatment (48 hours) with IC_50_ of the antibiotics. PC3 cells treated with puromycin show lesser cell migration compared to blasticidin-treated cells at 48 hours after treatment.

**Figure 6B:**
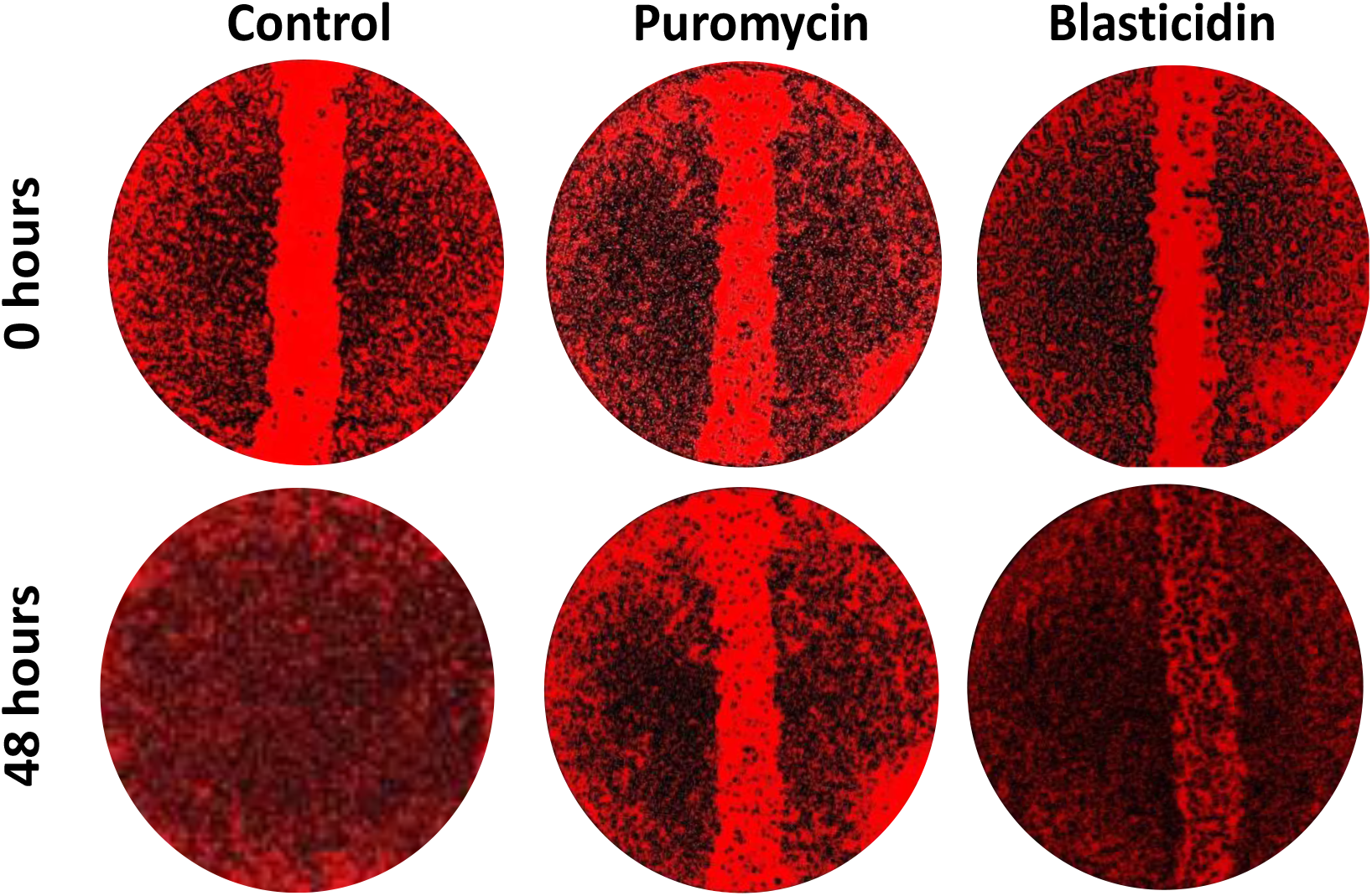
Photomicrograph of scratch cell-migration assay of DU145 colonies following treatment with puromycin and blasticidin antibiotics. Microscopic images (scratch cell-migration assay) showing colonies of DU145 before (0 hours) and after treatment (48 hours) with IC_50_ of the antibiotics. DU145 cells treated with puromycin show lesser cell migration compared to blasticidin-treated cells at 48 hours after treatment.

**Figure 6C:**
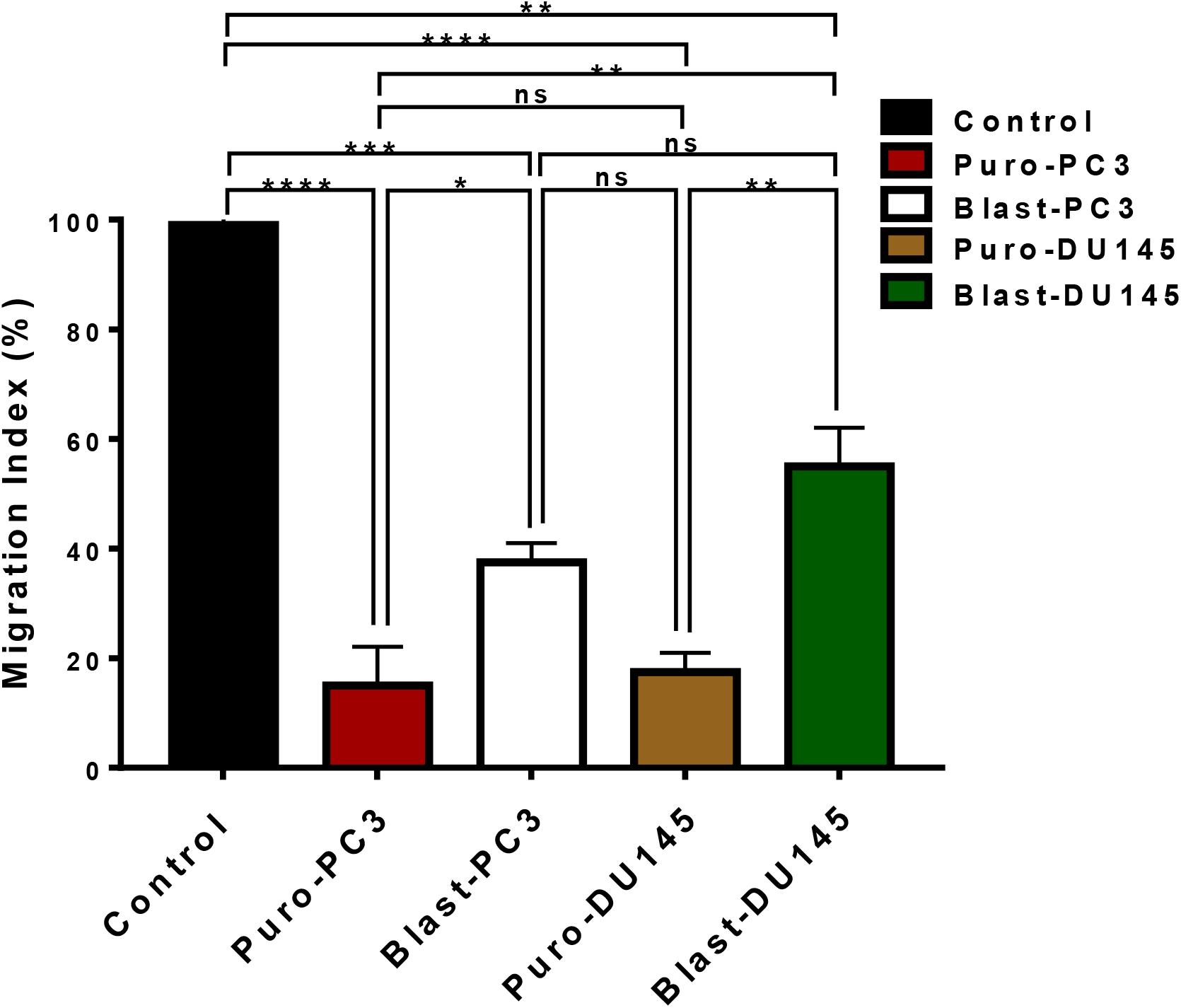
Bar chart comparing calculated migration indexes generated from scratch migration assay which compares the treatment-induced decrease in migration index between PC3 and DU145 cells following treatment (IC_50_) with puromycin and blasticidin relative to the control. *p < 0.05, **p < 0.01, ***p < 0.001, ****p < 0.0001 indicate significant p-values while ns indicates non-significant p-values. Treatments were done in triplicates and repeated at least 3 times independently.

### Puromycin and blasticidin antibiotics induced apoptotic cell death of mCRPC cells

At 48 hours post-treatment with the nucleoside antibiotics, we observed morphological changes in both PC3 and DU145 cell lines that are characteristic of the hallmarks of apoptosis. The untreated PC3 and DU145 cells looked healthy, spindle-shaped, and attached to surrounding cells, whereas the treated cells became detached from others, roundish, shrunken with a fragmented membrane, condensed and lysed nuclei (karyorrhexis), and apoptotic bodies, which are signs of apoptotic (programmed) cell death. These results reinforce the fact that puromycin is more effective in both PCa lines and that both antibiotics have an apoptotic effect on both cell lines.

These changes were found to be drug-dependent and dose-dependent. To confirm whether the mode of cell death induced by both antibiotics was truly due to apoptosis, we performed a dual fluorescent staining experiment using acridine orange (stains all cells green) and propidium iodide (stains dead cells yellowish-brown/red). In this study, we observed that both puromycin and blasticidin induced apoptotic cell death in two cell lines (**Figure 7A**). However, the degree of apoptotic staining relatively differs between both cell lines when exposed to either of the drugs (**Figure 7B**). When exposed to IC_50_ of puromycin or blasticidin, we observed more late-stage apoptotic (reddish-brown colored) cells in the PC3 cell line compared to the DU145 cell line, which contains mostly early-stage (yellow) apoptotic cells (**Figure 7A & 7B)**.

**Figure 7A:**
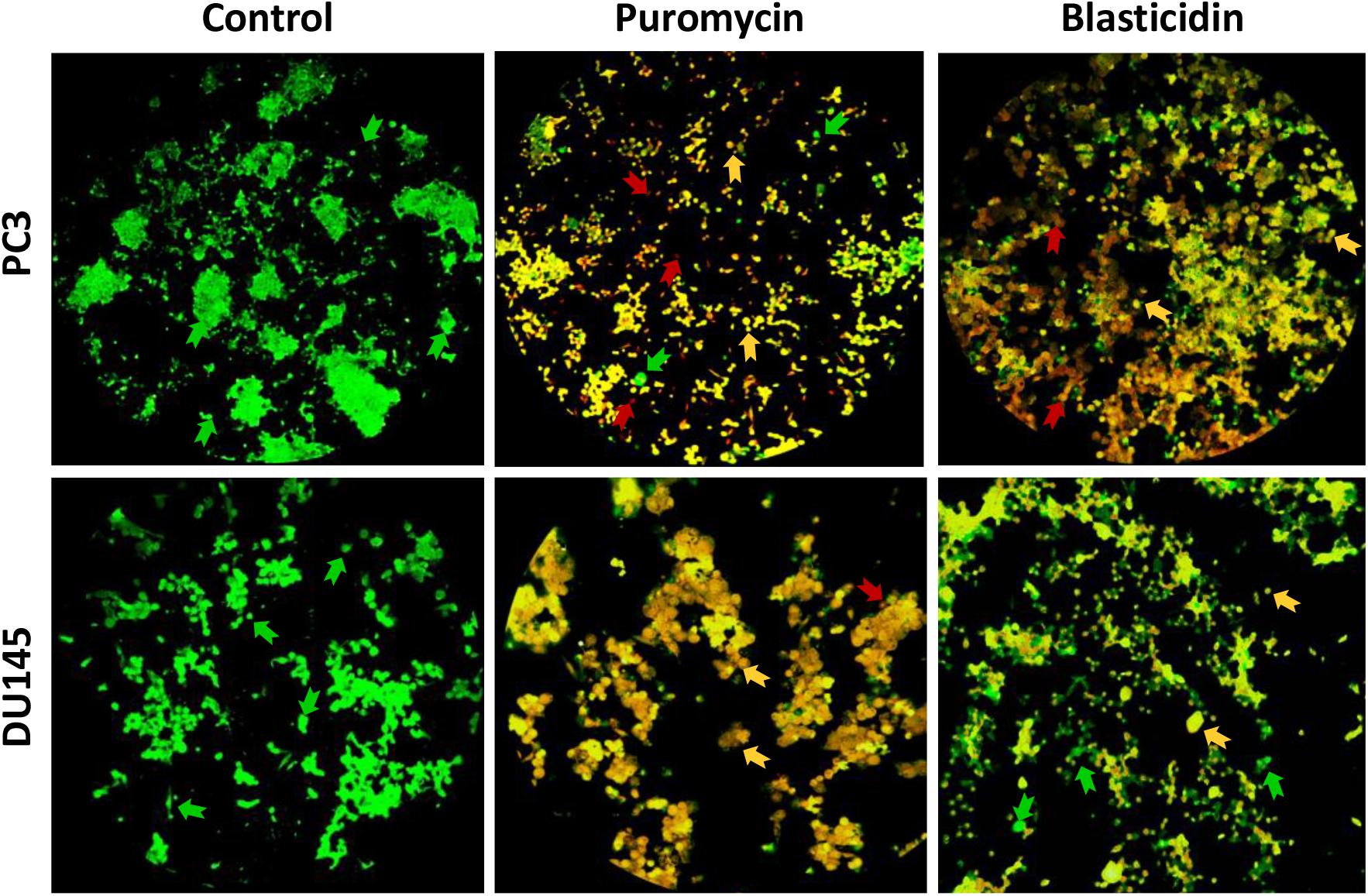
Photomicrographs of the dual immunofluorescent cell death assay of PC3 and DU145 cells after treatment with puromycin and blasticidin antibiotics. Microscopic images of acridine orange & propidium iodide dual-stained PC3 and DU145 cells following treatment with IC_50_ of puromycin and blasticidin for 48 hours. Acridine orange (AO) dye stains viable cells green while propidium-iodide (PI) dye stains apoptotic cells orange (early apoptosis) to red (late apoptosis). More puromycin-treated and blasticidin-treated PC3 cells were observed to be undergoing apoptosis compared to puromycin-treated and blasticidin-treated DU145 cells. The green, yellow and red arrows are pointing to live, early apoptotic, and late apoptotic cells, respectively.

**Figure 7B:**
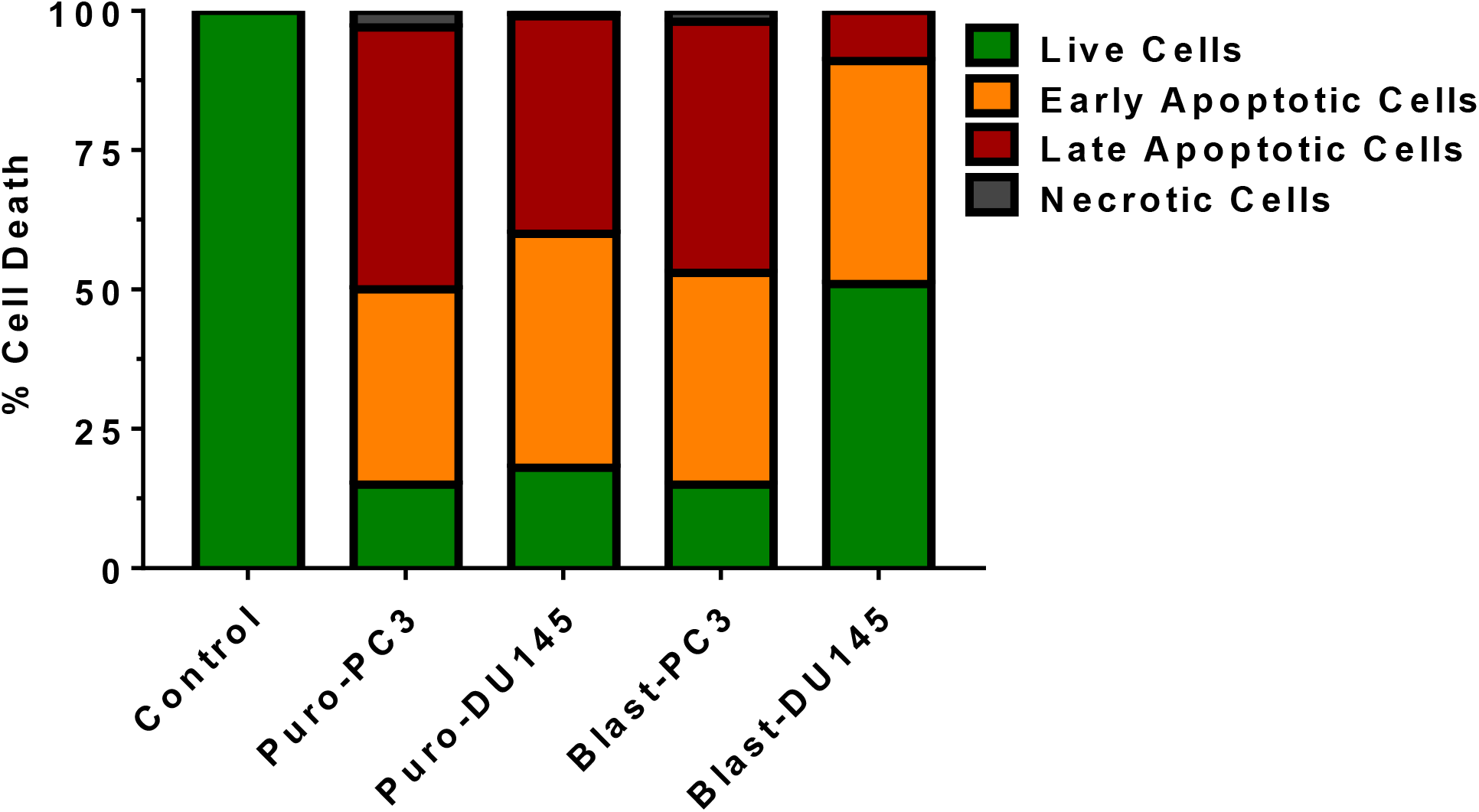
Bar chart showing the percentage of PC3 and DU145 cells at different stages of cell death compared to control after treatment with IC_50_ of puromycin and blasticidin. % of live cells is indicated by green bars, early apoptotic cells (%) in yellowish-orange bars, late apoptotic cells (%) in reddish-brown bars, necrotic cells (%) in black bars, Experiment was repeated at least 3 times independently.

### Puromycin and blasticidin antibiotics enhanced the anticancer effects of taxanes on mCRPCs

To evaluate the ability of both puromycin and blasticidin antibiotics to enhance the anticancer effects of taxanes in a combination treatment regimen, we treated PC3 cells with IC_50_ of either puromycin or blasticidin followed by treatment with graded doses (1 nM - 10 µM) of paclitaxel (PAC), docetaxel (DOC), and cabazitaxel (CAB). The IC_50_ values measure the amount of drug needed to exert half of its maximal inhibitory concentration. The higher the value the less efficient the substance is at disrupting the mechanism, while lower values indicate stronger inhibition. When comparing the results for paclitaxel treatment, we observed that the IC_50_ value decreased from 3 nM (PAC alone) to 0.9 nM when combined with puromycin, which is lower than the IC_50_ value of 1 nM measured when paclitaxel was combined with blasticidin. This indicates that puromycin is better at enhancing the anticancer effect (antiproliferative) of paclitaxel in PC3 cells than blasticidin (**Figures 8A & 8B**). Similar data patterns were observed for treatments involving docetaxel and cabazitaxel as well. Docetaxel decreased from 4.3 nM (DOC alone) to 0.6 nM when combined with puromycin and 0.8 nM when combined with blasticidin (**Figures 8C& 8D**). In addition, IC_50_ of cabazitaxel decreased from 0.7 nM (CAB alone) to 0.1 nM with puromycin and 0.2 nM with blasticidin (**Figures 8E & 8F**).

**Figure 8:**
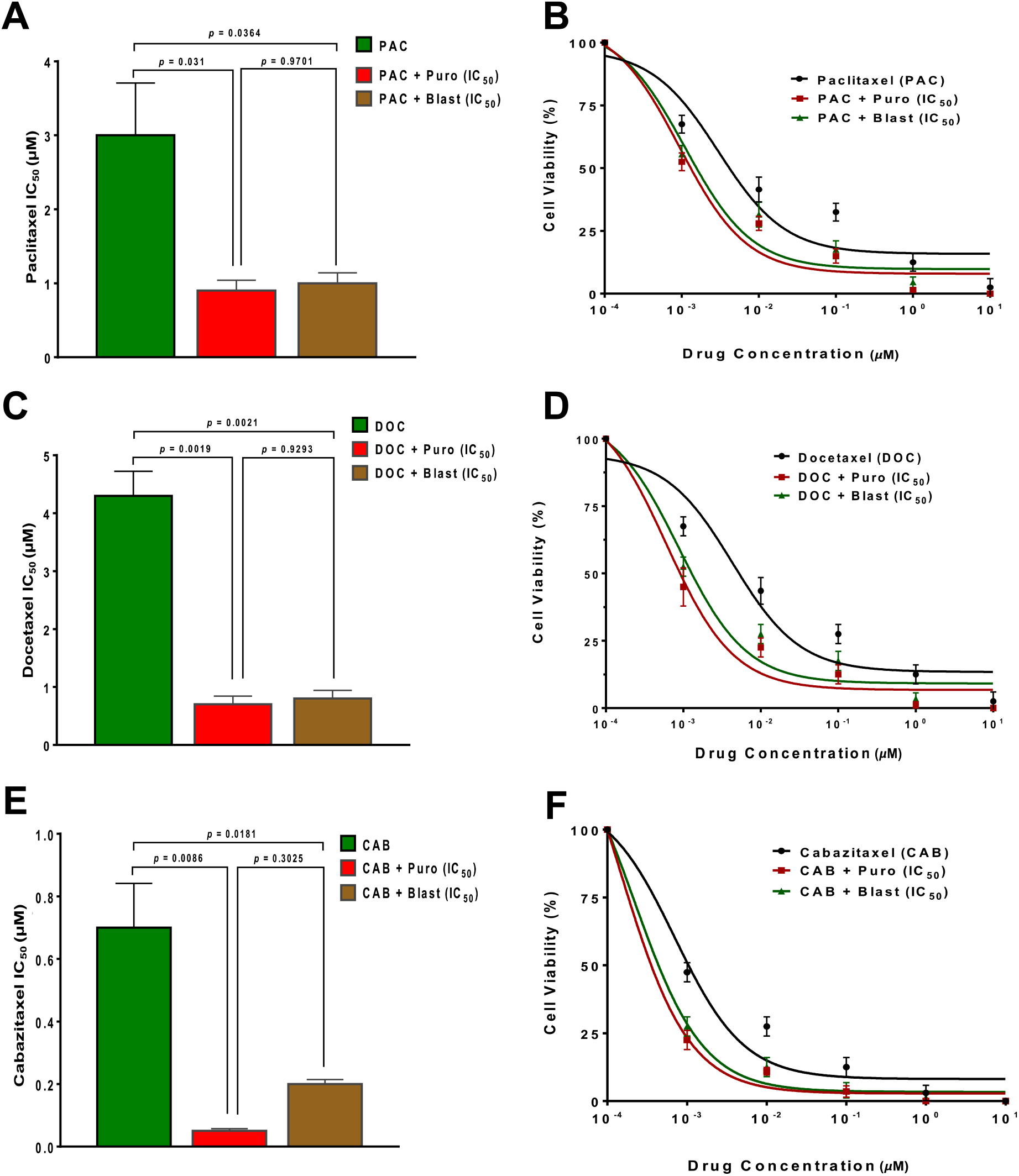
Dose-response plots showing cell viability of PC3 cells when treated alone with each taxane compared to when treated with a combination of each taxane with either puromycin (IC_50_) or blasticidin (IC_50_). **A & B**. Dose-response plot (non-linear regression fit) and IC_50_ bar plot of PC3 cells treated with paclitaxel (PAC) alone, PAC in combination with puromycin (Puro) (IC_50_), and PAC in combination with blasticidin (Blast) (IC_50_) for 48 hours. IC_50_ was determined to be 3 nM for PAC treatment alone, 0.9 nM for PAC plus Puro, and 1 nM for PAC plus Blast treatments. **C & D**. Dose-response plot (non-linear regression fit) and IC_50_ bar plot of PC3 cells treated with docetaxel (DOC) alone, DOC in combination with Puro (IC_50_), and DOC in combination with Blast (IC_50_) for 48 hours. IC_50_ was determined to be 4.3 nM for DOC treatment alone, 0.6 nM for DOC plus Puro, and 0.8 nM for DOC plus Blast treatments. **E & F**. Dose-response plot (non-linear regression fit) and IC_50_ bar plot of PC3 cells treated with cabazitaxel (CAB) alone, CAB in combination with Puro (IC_50_), and CAB in combination with Blast (IC_50_) for 48 hours. IC_50_ was determined to be 0.7 nM for CAB treatment alone, 0.1 nM for CAB plus Puro, and 0.2 nM for CAB plus Blast treatments. Doses of taxanes: 0, 0.01, 0.1, 1.0, 10 µM. Data are represented as mean ± SD in triplicates, in which we repeated the experiment independent 3 times. P < 0.05 is considered as significant.

To better assess whether the combination of the drugs was favorable or unfavorable, we calculated the IC_50_ dose-reduction index (DRI) which indicates the dose reduction between the combo and the single treatment at a given level (50% growth inhibition) in a synergistic combination. DRI was calculated using the formula DRI = D_single_ ÷ D_combo_, in which DRI = 1 indicates no dose reduction, DRI > 1 indicates favorable dose-reduction, and DRI < 1 indicates unfavorable dose reduction. The dose-reduction indices for paclitaxel, docetaxel, and cabazitaxel in combination with puromycin or blasticidin were determined to be favorable (DRI > 1). DRI of paclitaxel plus puromycin = 3.3, and DRI of paclitaxel plus blasticidin = 3.0. Also, the DRI of docetaxel plus puromycin = 7.2, and the DRI of docetaxel plus blasticidin = 5.4. While the DRI of cabazitaxel plus puromycin = 7, the DRI of cabazitaxel plus blasticidin = 3.5.

Another important therapeutic parameter assessed in our drug combination experiments was combination index (CI) at IC_50_, which is the change in effect between the combo and the single treatment at a given dose, at 50% efficacy. To calculate the CI, PC3 cells were treated with a combination of puromycin/blasticidin and paclitaxel/docetaxel/cabazitaxel using the method of constant ratio drug combination proposed by Chou and Talalay (Ting-Chao, 2010). CI was calculated using the formula; CI = (D1_Combo_ ÷ D1_Single_) + (D2_Combo_ ÷ D2_Single_), in which CI < 1 indicate synergistic effect due to drug combination, CI = 1 indicates addictive effect due to drug combination, and CI > 1 indicates antagonistic effect due to drug combination. D1_Combo_ and D2_Combo_ are the concentrations/doses of drug {1} and drug {2} used in combination to achieve 50% drug effect while D1_Single_ and D2_Single_ are the concentrations/doses for drugs administered alone to achieve the same effect. The combination indices for paclitaxel, docetaxel, and cabazitaxel in combination with puromycin or blasticidin were determined to be synergistic (CI < 1). CI of paclitaxel plus puromycin = 0.38, and CI of paclitaxel plus blasticidin = 0.44. Also, the CI of docetaxel plus puromycin = 0.14, and the CI of docetaxel plus blasticidin = 0.20. While the CI of cabazitaxel plus puromycin = 0.20, the CI of cabazitaxel plus blasticidin = 0.31.

## DISCUSSION

Antibiotic chemotherapy, otherwise known as antibiochemotherapy, involves the use of antibiotics with chemotherapeutic properties for the treatment of various cancers. In the past, several antibiotics with different mechanisms of action and pharmacokinetics have been investigated and found to have not just antimicrobial activities but also anticancer properties [**25**].

In this study, we hypothesized that both puromycin and blasticidin nucleoside antibiotics can inhibit mCRPC proliferation, metastasis, and induce cell death via apoptosis, in a dose- and time-dependent manner. This theory was inspired by previous studies demonstrating how nucleoside antibiotics and analogs such as gemcitabine, anthracycline, and epirubicin exert their anticancer effects when used in the treatment of PCa [**26**]. For instance, gemcitabine was shown to mediate its antitumor effects by promoting apoptosis of malignant cells by blocking the progression of cells through the G1/S-phase boundary and also found to be beneficial when administered with docetaxel for treatment of mCRPC [**27, 28**]. Furthermore, puromycin has both cytotoxic and antiproliferative effects, prompting us to investigate its ability to induce apoptosis and inhibit aggressive behaviors in mCRPCs [**29**].

Studies have also highlighted that blasticidin S (BlaS) is a potent inhibitor of protein synthesis in bacteria and eukaryotes, hence favoring its common use in genetic engineering [**30**]. BlaS is produced by the *Streptomyces species* and is a nucleoside analog consisting of a cytosine bonded to a pyranose ring and attached to an N-methyl-guanidine tail [**31**]. BlaS works by enhancing tRNA binding to the P site of the large ribosomal subunit and slowing down spontaneous intersubunit rotation in pre-translocation ribosomes [**32**]. Interestingly, BlaS can stabilize the deformed P-site of tRNA which in turn effectively inhibits peptidyl-tRNA hydrolysis by release factors, and, to a lesser extent, peptide bond formation [**33**]. At lower concentrations, BlaS was found to slow down peptide release by the activities of eRF1, eRF3a, and GTP, which partially prevents peptidyl-tRNA hydrolysis. Whereas at higher concentrations, BlaS inhibits peptide synthesis revealing an impact on both elongation and termination steps [**34**]. Because BlaS predominantly targets peptide release and also inhibits protein synthesis in eukaryotes, the anticancer properties of BlaS need further elucidation in various cancer cells [**35**].

Mutation, deletion, and deregulation of p53 have been described to be common in mCRPCs [**36**]. An increase in the expression of p53 in cells has been shown to induce a decrease in the proliferative index, and the survival rate of cancer cells [**37**]. This makes p53 a common target of investigation when studying PCa. Puromycin has been shown to induce p53-dependent apoptosis via upregulation of RPL5 or RPL11 for binding with MDM2 in human colon cancer cell lines [**38**]. With consideration to research showing puromycin’s ability to enhance p53 activity thereby causing antitumor effects, we investigated how puromycin and blasticidin behave in PC3 and DU145 cell lines with different p53 statuses [**39**]. Moreover, the PC3 cell line is known to be more aggressive, castration-resistant, and metastatic compared to the DU145 cell line, which is another moderately aggressive mCRPC cell line [**39, 40**].

One of the crucial aspects of our investigation is the antimetastatic effects of our antibiochemotherapy on mCRPCs. Metastasis is one of the hallmarks of PCa progression and is defined as the migration of cells from their tissue of origin to a distant or regional part of the body [**41, 42**]. The most common organs that PCa cells migrate to are the bones and lymph nodes, but they can also spread to the thorax, liver, and lungs, and less commonly to the kidneys, gastrointestinal tracts, and the retroperitoneum [**43, 44**]. To determine whether the nucleoside antibiotics (blasticidin and puromycin) can reduce the metastatic potentials of both mCRPC cell lines, we used the scratch migration assay to determine the antibiotic-induced effects on cell migration. Scratch migration assay is a process where a scratch is created at the center of the well plates and the diameters of the scratches are compared pre-treatment and post-treatment to measure the anti-metastatic effect of the drugs [**45**]. We observed the cell-dependent, drug-dependent, dose-dependent, and time-dependent anti-metastatic effects exhibited by both antibiotics. Puromycin shows a more significant antimetastatic activity in the two mCRPC cell lines compared to blasticidin. The anti-metastatic effects were found to be more potent in PC3 cells compared to DU145 cells. The reason for this is currently unknown but it could be due to the differences in aggressiveness and proliferative nature of both cell lines [**46**]. We extrapolate that both antibiotics inhibited PC3 cells at a quicker pace, due to their faster mitotic index compared to DU145 cells, which can be attributed to cell cycle arrest [**47**].

Since there have been cases linking PCa and prostatitis, physicians may prescribe antibiotics in anticipation of this event [**48**]. One of the most used antibiotics for chemotherapeutic treatment is doxorubicin, which has been shown to have great efficacy against a variety of cancers, such as PCa, leukemia, lymphoma, breast cancer, among others [**49**]. Gemcitabine is another example of nucleoside antibiotic analog that has been used either alone or in combination with other chemotherapies for the treatment of several types of cancer, including metastatic breast cancer, non-small cell lung carcinoma, ovarian cancer, PCa, and pancreatic cancer [**50**]. Being that both blasticidin and puromycin are effective against human and microbial cells makes them a befitting antibiochemotherapy for PCa [**51**]. For instance, antibiotics such as chloramphenicol and linezolid among others that are widely used bind to the A site of the large subunit of the ribosome, whereas blasticidin binds to the P site of the ribosomal unit where the peptidyl-transferase is being inhibited [**52**].

To observe the anti-tumorigenic effect of both nucleoside antibiotics, we performed a colony formation assay to measure colonies formed post-treatment in comparison to untreated cells [**53**]. This assay also serves as an indicator of the resistance status of the cells to the drugs or other agents of interest. Our results revealed that PC3 cell lines were better inhibited by puromycin than blasticidin. On the seventh day post-treatment, comparing the treated cells with the control for each cell line, we noticed a significant difference in the cell confluences and the number of colonies formed. Similarly, DU145 cells were better inhibited by puromycin than blasticidin, showing that DU145 cells may be less sensitive to blasticidin than PC3 cells. Before treatment, it is important to note that PC3 cells formed abundant colonies compared to DU145 cells, which showed sparsely distributed colonies. The reduced ability to form colonies after treatment alludes to the inability to form microtumors, which may be related to p53 activity in the cells as demonstrated in previous studies [**38, 54**].

It has also been reported that DU145 cells contain two mutations in the p53 gene: at codons 274 (proline to leucine) and 223 (valine to phenylalanine) [**55**]. Whereas PC3 cells contain a single p53 mutation (hemizygous) at chromosome 17p, in which codon 138 exhibits a base pair deletion causing a frameshift and in-frame stop codon at position 169. The fact that PC3 cells have only one p53 mutation while DU145 cells have two p53 mutations may contribute to the lower susceptibility of DU145 cells to the antiproliferative effects of p53 mediated by puromycin [**56**]. This is possibly one of the major reasons for the disparity in the response of both mCRPC cells to the nucleoside antibiochemotherapy [**57**].

We also found from our results that the effect of both nucleoside antibiotics on the colony-forming properties of the PCa cell lines was dose-dependent, in which for every increase in the antibiotic concentration, there were fewer colonies formed. Additionally, it seems treatment with IC_50_ of antibiotics effectively inhibits colony formation in PC3 cells rather than DU145 cells. We also observed that these effects are time-dependent; meaning that the longer we expose the cells to the antibiotics, the higher the decrease in cell proliferation, survival, or growth.

The MTT and Resazurin assays allowed us to observe the effects of blasticidin and puromycin on PCa cell viability and proliferation [**58**]. The results demonstrate a cell-dependent, drug-dependent, and dose-dependent response. The MTT reduction assay highlights the activity and quality of the mitochondrial contents. Cell viability is positively correlated to mitochondrial functionality. Consequently, the better functioning the mitochondria, the more it can actively reduce MTT into formazan by converting NADH to NAD+ [**59**]. Resazurin utilizes a similar mechanism as MTT but instead, resazurin is reduced to resorufin [**60**]. Conversely, when the mitochondria are not functioning properly they cannot provide the necessary energy for the cells to actively metabolize nutrients, resulting in reduced cell proliferation, viability, and a reduction in the assay color change when observing the well plates [**61**]. PC3 cells were more susceptible when exposed to blasticidin and puromycin in comparison to DU145 cells. One explanation to consider may be the fact that PC3 cells undergo a faster rate of cell cycle/division (shorter doubling time) and therefore proliferate at a higher rate than DU145 cells (longer doubling time) [**39, 54**]. This increased rate of proliferation means increased demand for energy and cell signaling allows, which may be inhibited by nucleoside antibiotics [**40**].

Also, nucleoside antibiotics enter cells through specific nucleoside transporters. The cellular uptake of nucleoside antibiotics is an active process that involves the service of concentrative nucleoside transporters (CNTs) and equilibrative nucleoside transporters (ENTs) [**62**]. About three CNTs and four ENTs have been identified and described in humans. The number of nucleoside transporters on the cancer cell membrane may also dictate their degree of susceptibility to the nucleoside antibiochemotherapy.

As displayed in our results, puromycin requires a lower concentration than blasticidin to exhibit antiproliferative effects on PCa cells, despite both antibiotics having similar binding sites which indicate that they may have similar functionality [**63**]. Some factors that may reduce the activity of blasticidin include an organism’s ability to generate BS deaminase and acetylation, as well as the increased abundance of nucleoside transporters [**64**]. BSD and BSR are two known BS deaminase genes. BS deaminase’s enzymatic process results in the conversion of cytosine to 4-diamino-4-hydroxy blasticidin in BSD [**65**]. Furthermore, our results indicate a possible relationship between cell viability/proliferation and the dose of antibiotics given. As the antibiotic dosage increase, cell viability and proliferation decrease.

To further determine whether treatment with either nucleoside antibiotics will induce apoptotic cell deaths, we used acridine orange and propidium iodide dyes to stain the mCRPC cells after treatment with antibiotics to identify the mode of cell deaths. Acridine orange stains only live cells, whereas propidium iodide stains dying cells (nucleic acids) with a damaged cell membrane [**66**]. Observing the different ranges of fluorescent colors from yellow (early-stage) and brown-red (late-stage) will reveal the different stages of apoptosis. Cell deaths, caused by puromycin and blasticidin in our experiment, occurs by two different pathways; the preferable one will be apoptosis versus necrosis [**67**]. The mechanism used by apoptosis can occur through the intrinsic pathway where a cell undergoes programmed cell death due to stress, or the extrinsic pathway where a cell undergoes apoptosis due to signals received from other cells [**68**]. On the other hand, necrosis is another mode of chemotherapy-induced cell death, in which expulsion of the cell cytosolic component leads to inflammation and damages to neighboring normal cells [**69**]. In our study, we found that both nucleoside antibiotics induced apoptotic cell deaths, which is therapeutically preferable. The marked apoptosis observed after nucleoside antibiochemotherapy suggests a stress-induced response in the mCRPC cells, in which blasticidin or puromycin may have activated apoptosis-mediated pathways. The reactivation of p53 protein-mediated cell cycle arrest and p53-mediated apoptosis of PCa cells is another yet to be explored mechanism of cell death [**54, 70**].

Due to the problematic nature of drug toxicity at high doses, scientists and clinicians in the past have tried combining two or more drugs synergistically at lower doses [**71**]. The reduction in the inhibitory dose (IC_50_) of each taxane by puromycin or blasticidin is important in decreasing drug-associated side effects and the expensive costs associated with taxanes. We believe that administering a lower concentration of each drug in a synergistic combination regimen will provide a robust anticancer strategy for the elimination of cancer cells in prostate cancer patients, as indicated by the combination of nucleoside antibiotics and taxanes in our study.

Also, the fact that prolonged overall survival (OS) and progression-free survival (PFS) has been reported in pancreatic cancer patients who received gemcitabine-based chemotherapy as first-line therapy, compared to those receiving 5FU-based chemotherapy further provides a proof of concept of the potential benefit of nucleoside antibiotic usage in patients with mCRPC when administered alone or in combination with other drugs [**72**]. Since there are currently no definitive curable treatment options for mCRPC, more breakthrough therapeutic combination options that incorporate antibiotics may be needed to effectively treat mCRPCs in men [**73**].

## CONCLUSION

We have been able to establish the therapeutic potential of puromycin and blasticidin in the treatment of mCRPCs. Further studies would focus on understanding the molecular mechanisms of both antibiotics in mCRPCs and the validation of these findings in animal models. Other focus will be geared towards developing strategies to limit the side-effects and off-target effects of treatment with antibiotic compounds, including their packaging in functionalized and biodegradable nanoparticles that select for cancer cells only. Other strategies involve identifying highly selective and efficacious nucleoside analogs with fewer side effects. In conclusion, our study shows that both puromycin and blasticidin nucleoside antibiotics can induce apoptotic cell deaths and inhibit cell migration and proliferation of mCRPCs, thus justifying the need for more molecular studies to understand the mechanisms of their anticancer effects in mCRPCs.

## AUTHOR CONTRIBUTION

All primary authors contributed equally to the execution of this project and the writing, reviewing, and editing of the manuscript before submission.

## ACKNOWLEDGEMENT

This research project was partially funded by a grant awarded to G. A. L, F.A, M. P, and J. B. from the Office of Undergraduate Research and Inquiry (OURI). We are also grateful to the Dean’s Office of FAU Charles E. Schmidt College of Science for the supplemental funding awarded to S. O. O through the Science Graduate Research Scholarship and Vincent Saurino Fellowship.

